# Increased error rate and delayed response to negative emotional stimuli in antisaccade task in obsessive-compulsive disorder

**DOI:** 10.1101/2023.06.21.545955

**Authors:** Guzal Khayrullina, Olga Martynova, Elizaveta Panfilova

**Affiliations:** Institute of Higher Nervous Activity and Neurophysiology RAS, Butlerova 5A, Moscow, 117484, Russia; Centre for Cognition and decision making, Institute for Cognitive Neuroscience, National Research University Higher School of Economics, Myasnitskaya 20, Moscow, 101000, Russia

**Keywords:** obsessive-compulsive disorder, eye-tracking, inhibitory control, attention, emotion.

## Abstract

Ample evidence links impaired inhibitory control, attentional distortions, emotional dysregulation, and obsessive-compulsive disorder (OCD). However, it remains unclear what underlies the deficit that triggers the OCD cycle. The present study used an antisaccade paradigm with emotional stimuli to compare eye movement patterns reflecting inhibitory control and attention switching in OCD and healthy control groups. Thirty-two patients with OCD and thirty healthy controls performed the antisaccade task with neutral, positive, and negative visual images. The groups differed significantly in the number of errors to negative stimuli. The latency of correct antisaccades varied between valences in OCD group when negative and neutral stimuli served as target ones. The OCD group showed more errors to negative stimuli than the control group and they also performed antisaccades more slowly to negative and neutral stimuli than positive ones. Other patterns, including mean velocity of correct antisaccades and anticipatory saccades, did not differ between groups. The mean velocity of correct antisaccades was higher for negative and positive stimuli than for neutral stimuli; however, there were more anticipatory saccades for neutral stimuli than for negative and positive stimuli in both groups. The peak velocity parameter did not show any differences either between groups or between valences. The findings support a hypothesis that an attentional bias towards threatening stimuli interferes with inhibitory control in OCD.

## 1. Introduction

**Obsessive-compulsive disorder (**OCD) has been named one of the top ten disabling disorders by the World Health Organization (WHO, 1999). The ICD-10, ICD-11 and DSM-5 describe OCD as the presence of repetitive obsessive thoughts or compulsive actions. Obsessive thoughts are ideas, images, or impulses that enter an individual’s mind over and over again in a stereotyped form that the person cannot get rid of. However, patients recognize those thoughts as his or her own, even if they are involuntary and often disgusting. Compulsions or rituals are stereotyped behaviors that are repeated over and over again. They are not inherently enjoyable and do not lead to inherently useful tasks. Their function is to escape from the anxiety state, connected with preventing some objectively unlikely event (APA, 2013; WHO, 1983, 2022).

Clinically, the symptoms of OCD are manifested in both the inability to stop negative repetitive thoughts and switch them to more productive ones and to prevent the performance of useless actions. In connection with the clinical symptoms, neuropsychological studies have asserted the existence of cognitive deficits in this patient category, including disruptions of inhibitory control (Benzina et al., 2016). Impairment of inhibitory control is one of the most central features of OCD (Chamberlain et al., 2005). However, the results of numerous neuropsychological studies are inconsistent (Blom et al., 2011; Gruner & Pittenger, 2017; Lipszyc & Schachar, 2010; Marincowitz et al., 2022; Morein-Zamir et al., 2013). This may be due to the fact that different models of tasks investigating the specific features of inhibitory control were used in the studies, and there was no distinction between inhibitory control into the cognitive and behavioral aspects of inhibition. These factors could have influenced the results of the studies (Benzina et al., 2016). However, meta-analyses of OCD have shown convincing data on the impairment of both the cognitive and behavioral profile of inhibitory control (Abramovitch et al., 2013; Shin et al., 2014; Snyder et al., 2015).

Emotional states also affect inhibitory control and may cause impairment (Bannon et al., 2008; Bohne et al., 2005; Zetsche et al., 2015). In the study of (Bannon et al., 2008), participants with OCD showed reduced inhibitory control in response to threatening negative pictures. It can be explained with a conception that cognitive theories emphasize the fact that manifestations of negative emotions are associated with rigidity in information processing rooted from the individual’s preoccupation and response to perceived danger (Beck A.T. et al., 1985; Williams J. M. et al., 1997). People with OCD have rigid ideas about the personal meaning of these thoughts and their catastrophic consequences, which lead to high levels of anxiety, distress, or guilt (Rachman S. & Hodgson R., 1980). Impaired cognitive profile of inhibitory control in patients with OCD manifests under emotionally negative circumstances (Morein-Zamir et al., 2013). Moreover, OCD patients showed deficits in conditioned fear extinction (Milad et al., 2013), which may indicate that the negative emotional stimulus is the primary factor influencing slow inhibition.

Neuroimaging data accumulated since the 1980s has led us to a biological hypothesis about the pathophysiology of OCD. The altered orbitofrontal subcortical circuit (OCD-loop) model can lead to the development of OCD symptoms and may be the basis for explaining the OCD phenomenon (Karpinski et al., 2017; Saxena et al., 1998). However, another review noted structural changes and altered functional activity of the limbic areas (including the amygdala), parietal and occipital cortex, and cerebellum (Hazari et al., 2019).

The model of Mataix-Cols et al. suggests that the different manifestations of obsessive-compulsive symptoms are mediated by relatively different components of the fronto-striatal-thalamic circuits involved in cognitive and emotional processing and that OCD can best be thought of as a spectrum of multiple, potentially overlapping syndromes rather than a single entity (Mataix-Cols et al., 2004). Symptom provocation tasks and emotion processing tasks have elicited the amygdala activation in OCD (Cardoner et al., 2011; Simon et al., 2014), with the highest correlation between aggression/checking and sexual/religious symptoms parameters. The increased amygdala activation in OCD appears to be specifically modulated by the type of symptom. The origin of such activation may be more closely related to the putative amygdala-centric pathway associated with abnormal fear processing (Via et al., 2014).

Impaired inhibitory control in patients with OCD has also been shown by eye-tracking method, especially using antisaccade task (Khayrullina et al., 2022). Antisaccade task is a reliable and sensitive tool in psychopathology, in particular concerning OCD, because the rate of success in antisaccade task depends on the integrity of the fronto-subcortical regions of the brain (Hutton & Ettinger, 2006; Narayanaswamy et al., 2021). In antisaccade task the subjects must suppress the reflex eye movement (prosaccade) to the visual stimulus and make a voluntary movement to the opposite point in the visual space. An erroneous reaction is an inability to suppress the reflex reaction of eye movements to a peripheral stimulus. These errors are usually followed by a corrective saccade indicating that the instruction was understood but the reflex response could not be suppressed (Luna et al., 2008).

Studies investigating inhibitory control with the antisaccade task have found increased error rates (Agam et al., 2014; Narayanaswamy et al., 2021) and increased latency of correct antisaccades in OCD patients (Maruff et al., 1999; Van Der Wee et al., 2006) compared to controls. The first evidence for an OCD endophenotype came from the Lennertz et al. study, which showed that OCD patients and their healthy first-degree relatives had an increased error rate and increased latency in the antisaccade task compared to healthy controls (Lennertz et al., 2012). Afterward, Bey K. supported this fact in a fairly large sample (169 patients with OCD), revealing increased antisaccade latency, intrasubject variability in antisaccade latency, and an increase in antisaccade errors compared to healthy controls. The latter effect was due to errors in express saccades, with OCD patients not significantly different from controls in terms of errors in regular saccades. Healthy relatives of OCD patients also showed an increased error rate and increased variability of the latent period in antisaccade tasks (Bey et al., 2018). Thus, most studies of antisaccades point to impaired inhibitory control in patients with OCD, manifested as an increase in latency and error rate which possibly correlates with an imbalance in the transmission of excitation and inhibition in the cortico-striatal-thalamo-cortical circuit. However, all of these studies used neutral stimuli like dots or circles.

Various studies confirm the influence of emotional stimuli on eye-movement patterns in OCD patients (Armstrong et al., 2010, 2012; Basel et al., 2023; Bradley et al., 2016). Armstrong et al. investigated features of attentional biases to threatening facial pictures, in which they determined increased vigilance for threat during free viewing and visual search but difficulty of stopping to threat in visual search tasks in OCD patients (Armstrong et al., 2012). Moreover, the group with severe OCD symptoms oriented attention to fearful faces and disgusted faces more than the group with mild OCD (Armstrong et al., 2010). The severity of OCD symptoms correlated with greater frequency and duration of fixation on OCD-relevant stimuli, which possibly reflected an attentional maintenance bias (Bradley et al., 2016). All of these findings consistently showed increased attention biases to negative and OCD-related pictures. However, there is no data on the influence of emotional stimuli on inhibitory control in OCD.

This study aimed to investigate the neurophysiological correlates of cognitive and emotional control in OCD using the antisaccade task. In particularly, the study compared eye-movement patterns (error rate, mean latency of correct antisaccades, mean velocity of correct antisaccades, peak velocity of correct saccades, anticipatory antisaccades) upon presentation to neutral, positive, and negative stimuli in the control group and the OCD group. The hypothesis of our study was that the parameters of oculomotor reactions between the control and OCD groups should differ more in response to negative stimuli than to positive and neutral stimuli. We used an overlap-type antisaccade task with fixation stimuli of different emotional valences (neutral, positive, and negative), where the participant first had to look at the central fixation stimulus surrounded by squares. After the disappearance of the fixation stimulus, a square remained on one of the sides; the participant had to look in the opposite direction from the square.

## 2. Methods

### 2.1. Participants

Thirty-two OCD patients (24.5 +/- 6.17 years old, 21 females) and thirty healthy volunteers (22.5 +/- 5.15 years, 17 females) participated in the study. Participants were all right-handed. A structured interview with patients was conducted by a clinical psychologist (G.K.) to ensure that patients with OCD met the criteria of the International Statistical Classification of Diseases and Related Health Problems 10th Revision (Brämer, 1988). 23 patients from the OCD groups received medication therapy but 9 participants did not (Table 1).

**Table 1.** Medications taken by members of the OCD group.

| Specification of the medication | Number of participants |
| --- | --- |
| selective serotonin reuptake inhibitors (antidepressants) | 11 |
| selective serotonin reuptake inhibitors (antidepressants) + antipsychotics | 4 |
| selective serotonin reuptake inhibitors (antidepressants) + anticonvulsants | 4 |
| selective serotonin reuptake inhibitors (antidepressants) + benzodiazepine | 1 |
| antipsychotics + anticonvulsants | 3 |
| without medication | 9 |

All patients were in a stable phase of the disorder during participation in the study. Individuals with drug abuse, neurological or mental disorders other than OCD (bipolar disorder, autism spectrum disorder, schizophrenia) were excluded from the group.

Healthy controls were recruited from volunteers via the announcement on social media. All the participants passed the assessment with the clinical psychologist (G.K.). The exclusion criteria were the following: mental and neurological disorders, brain trauma, and uncorrected vision.

### 2.2. Ethical statement

All experimental procedures complied with the requirements of the Helsinki Declaration. The ethical committee of the Institute of Higher Nervous Activity and Neurophysiology of the Russian Academy of Sciences approved the study protocol (#0125022021). All participants gave written informed consent before they participated in the study. Data collected from the participants were anonymized and concealed.

### 2.3. Psychological assessment

Participants filled in web-forms with questionnaires based on Russian versions of the Yale-Brown Obsessive Compulsive Scale (Y-BOCS).

The Yale-Brown Obsessive Compulsive Scale (Y-BOCS) is the scale developed by Wayne K. Goodman and his colleagues for the assessment of OCD severity (Goodman, 1989a). The total score consists of 10 core items divided by the subscales for obsessions (items 1–5) and compulsions (items 6–10) (Kim et al., 1994). Each item is evaluated on a 5-point system from 0 to 4 points. The assessment of the total score includes the following parameters: 0-7 - subclinical condition; 8-15 - an obsessive-compulsive disorder of mild severity; 16-23 - an obsessive-compulsive disorder of moderate severity; 24-31 - a severe obsessive-compulsive disorder; 32-40 - an obsessive-compulsive disorder of extremely severe severity. The scale is widely used in clinical practice and for scientific purposes, having high validity and reliability (Goodman, 1989a, 1989b; Rosario-Campos et al., 2006). Table 2 contains the group scores of levels of OCD symptoms according to Y-BOCS and the mean age for both groups.

**Table 2.** The group descriptive statistics.

| Test Score | OCD | Control group |
| --- | --- | --- |
| Yale-Brown Obsessive Compulsive Scale (Y-BOCS) | 20.9±8.27 | 1.2±2.06 |
| Mean age | 24.5±6.17 | 22.5±5.15 |

\*values represent means and standard deviations

### 2.4. Procedure and eye-movement data acquisition

Two days before the experimental procedure, participants obtained the preliminary screening tests before inclusion in the study in electronic form. Upon arrival at the research facilities, participants signed informed consent and consent to the processing of depersonalized data. After that, a clinical psychologist (G.K.) examined the participants using the structured clinical interview and later they fulfilled the Y-BOCS. The study was carried out using the equipment of the Research Resource Center # 40606 of IHNA and NPh RAS ‘Functional Brain Mapping’. The experiment was held in an eye-tracking laboratory equipped with a soundproofing dark room to keep consistent lighting conditions. Each participant got acquainted with the laboratory and passed the test version of the paradigm. Subsequently, the participants were asked to sit in a chair in front of a computer monitor.

Before the eye tracking experiments, ocular dominance was assessed using the hole-in-the-card test (Dolman method) (Cheng et al., 2004). In this test, the participant was instructed to hold a piece of cardboard with a central circular hole through which they had to view a target at about 6 m away with both eyes open. Subsequently, each eye was occluded in turn. The target would not be seen through the hole when the dominant eye was covered; on the contrary, the target persisted to be seen when the non-dominant eye was covered since the dominant eye would continue to fix the target. In this forced-choice test of dominance, there was only one result for dominance (left or right). The eye movement data were recorded using the dominant eye to avoid the potential confounding effect of differential dominance on eye tracking measures (Vergilino-Perez et al., 2012).

Eye-movement data were recorded by the eye tracker Eyelink Portable Duo (Sr Research Ltd., Canada) with a sampling rate of 500 Hz. The participant’s chin was comfortably fixed on a head mount to ensure stability. The 20″ flat screen monitor (Asus Vision XG248q, 240 Hz) had a resolution of 1152 × 864 pixels and was positioned 70 cm from the participant. All participants have got detailed instructions on how to perform a task. During the experiment, participants were asked to try to keep their heads as still as possible. Calibration and validation procedures were performed immediately before the task block. The participants were asked to visually follow a white dot moving in different places on the screen 9 times. The calibration time was about 30 seconds. Validation was carried out according to the same technical principles as calibration. If the accuracy was poor (fewer than 0.5**°**), recalibration was performed. The experiment was only started if the participant successfully passed the calibration and validation procedures. After that, the task instruction appeared on the screen depending on each block. After completing each block, the participant could rest for about 5 minutes. Before proceeding with the next block, the participant passed again the calibration and validation procedure. Overall, it took participants about 30-35 minutes to complete the experiment.

### 2.5. Antisaccade task

The paradigm was set using Eyelink Experiment Builder 2.3.1 software (Mississauga, Ontario, Canada: SR Research Ltd., 2020). The paradigm consisted of three blocks of antisaccade tasks (Subramaniam et al., 2018; Taylor and Hutton, 2009) with overlap-design. Each participant performed a total of 300 antisaccade trials in all three blocks. Each block contains 100 trials, of which 60 trials are pictures of neutral valence, 20 trials of positive valence, and 20 trials of negative valence. All of them were fixation stimuli that were taken from the International Affective Picture System (Lang P. J. et al., 2008). The fixation pictures were squares 250*300 mm, and the target stimuli were small squares 14*14 mm which were on both sides of the fixation point. Small squares were at ±12 degrees of visual angle from the center of the fixation picture.

The presented positive stimuli had a mean valence of 7.40 ranged 7.0–7.8, and a mean arousal of 4.9 ranged 4.2–5.6. The selected negative pictures had a mean valence of 2.2 (1.7–2.7) and a mean arousal of 6.1 (5.4–6.8). A circle serves as the neutral stimulus, which did not change in all presentation blocks. The angle of view on the stimulus was 41 degrees. The stimuli for each trial appeared on a screen with a black background.

Figure 1 illustrates the design of the experimental paradigm. Each block consisted of the antisaccade task with the Overlap design. The fixation stimulus of blocks comprised the pictures (neutral, positive, negative). Each trial began with a central fixation stimulus, which remained on screen for between 700-1500 ms. On both sides of the central fixation stimulus, there were 2 small squares. After an interval (700 to 1500 ms; interval occurring randomly, - in order to prevent the participants from the additive effect, which could affect the results), only the fixation stimulus with one square on one side remained on the screen, and the second square disappeared. After 200 ms the central fixation stimulus disappeared while the square was left on one side of the screen. The target stimulus (the square) lasted for 800 ms at two possible locations, +/-6° of visual angle from the center. The instruction to the individuals for the task was to look at the mirror image location of the remaining square. After the participant performed the task, a black screen appeared in blocks (break between trials) during 1000 ms (Figure 1). All blocks had a different set of negative and positive pictures.

**Figure 1.**
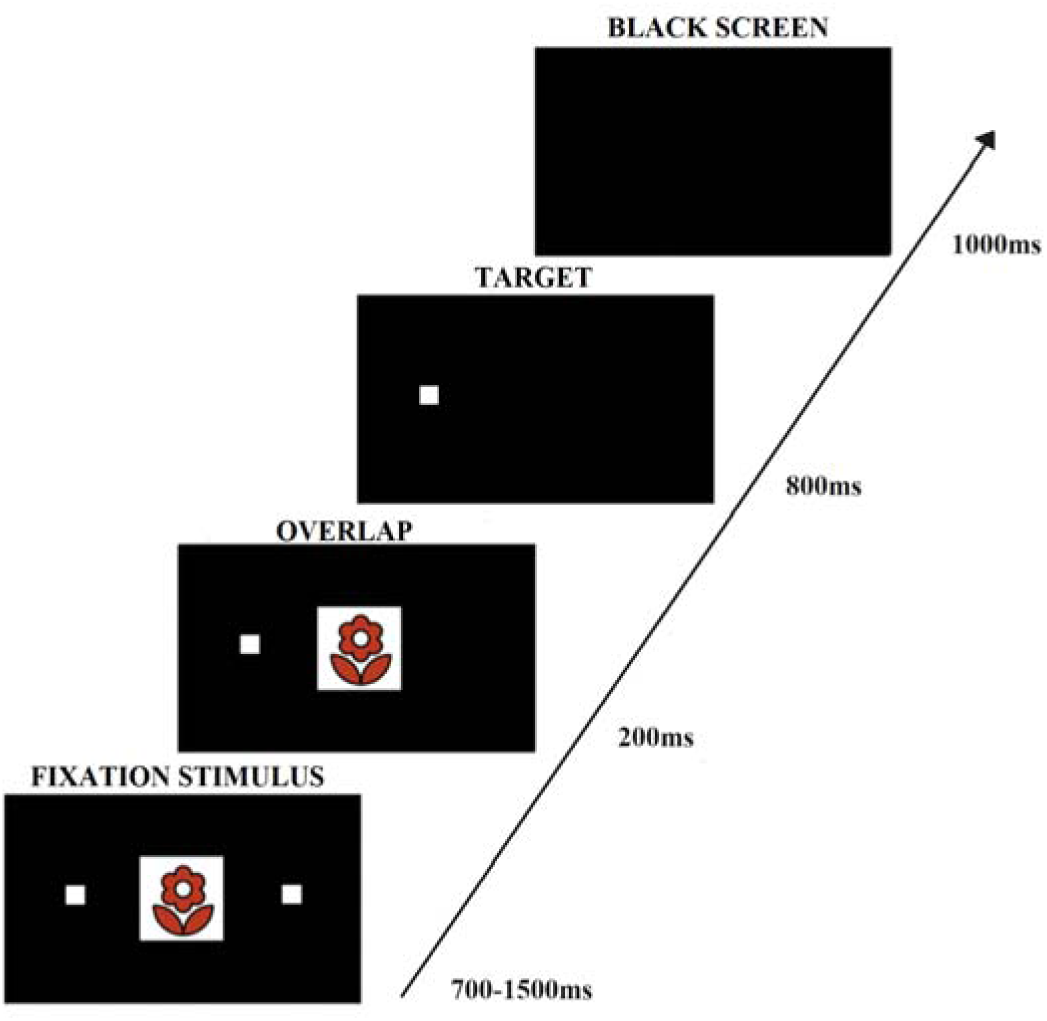
The Overlap design of antisaccade task paradigm with a substitute of stimulus the stimulus.

### 2.6. Data analysis

Data preprocessing was conducted using Data Viewer (SR Research). Trials with artifacts (blinks, etc.) were excluded from the analysis. Further analysis was performed in R Studio (https://www.rstudio.com/).

The following parameters were measured for all participants in both groups for each type of emotional stimulus: error rate; mean correct antisaccade latency; mean correct antisaccade velocity; mean correct antisaccade peak velocity; anticipatory saccade rate.

Parameters (error rate, mean correct antisaccade latency, mean correct antisaccade velocity, mean correct antisaccade peak velocity) were calculated with latencies more than 80 ms. Error eye directions were taken as errors if the person did not look in the mirror image from the presented target stimulus, including when the person looked at the stimulus, and also “up” and “down”. Trials with latencies less than 80ms were considered anticipatory saccades and multiplied by 100%.

### 2.7. Statistical analysis

The Shapiro–Wilk test was used to verify the normal distribution of samples. F-test was applied to compare the variances of two samples from normal distributions. Depending on the normality of distribution between-group age and self-reporting test difference was compared by Mann-Whitney test (for Y-BOCS, age difference) between the groups. The chi-square test checked the uniformity of sex distribution across groups. Statistical analysis of eye tracking measures was conducted in both groups for each parameter for each type of emotional stimulus. The mixed ANOVA design used 2 levels of between-group comparison (healthy controls versus participants with OCD) and 3 levels within-group comparison of the stimuli valences (neutral, positive and negative). If the ANOVA showed a significant effect, the Bonferroni post hoc method was used to make pairwise comparisons. Only results of statistical tests passed the p-value threshold of 0.05 are reported.

## 3. Results

The groups (healthy controls vs. OCD) did not differ significantly in the age distribution (22.5 ± 5.2 vs. 24.5 ± 6.2 years; p > 0.05). Gender distribution was also equal between groups (X-squared = 0.52019, p-value = 0.4708).

The level of obsessive-compulsive symptoms (Y-BOCS – 20.9±8.27 vs. 1.2±2.06, p>0,05) significantly differed between OCD and healthy control groups.

Eye-movement patterns significantly showed differences between groups in error rate (p<0.01). Latency differed markedly within the OCD group in negative (p<0.01) and neutral (p<0.05) stimuli. Mean correct antisaccade velocity (p<0.01), and anticipatory saccades (p<0.01) revealed differences in groups - only between valences. Peak velocity did not differ between groups and within groups between valences.

The error rate (Fig.2) during antisaccade performance varied significantly between groups in negative stimuli (p<0.01). The error rates for negative stimuli (p<0.01) differed markedly between groups, and the OCD group showed a higher error rate in negative stimuli. Intergroup analysis revealed statistically significant differences in the error rate between negative and neutral (p<0.01), negative and positive stimuli (p <0.01) in the OCD group. In the healthy control group, statistical differences in the error rate were not found between stimuli. Means and standard deviations are given in Table 3.

**Figure 2.**
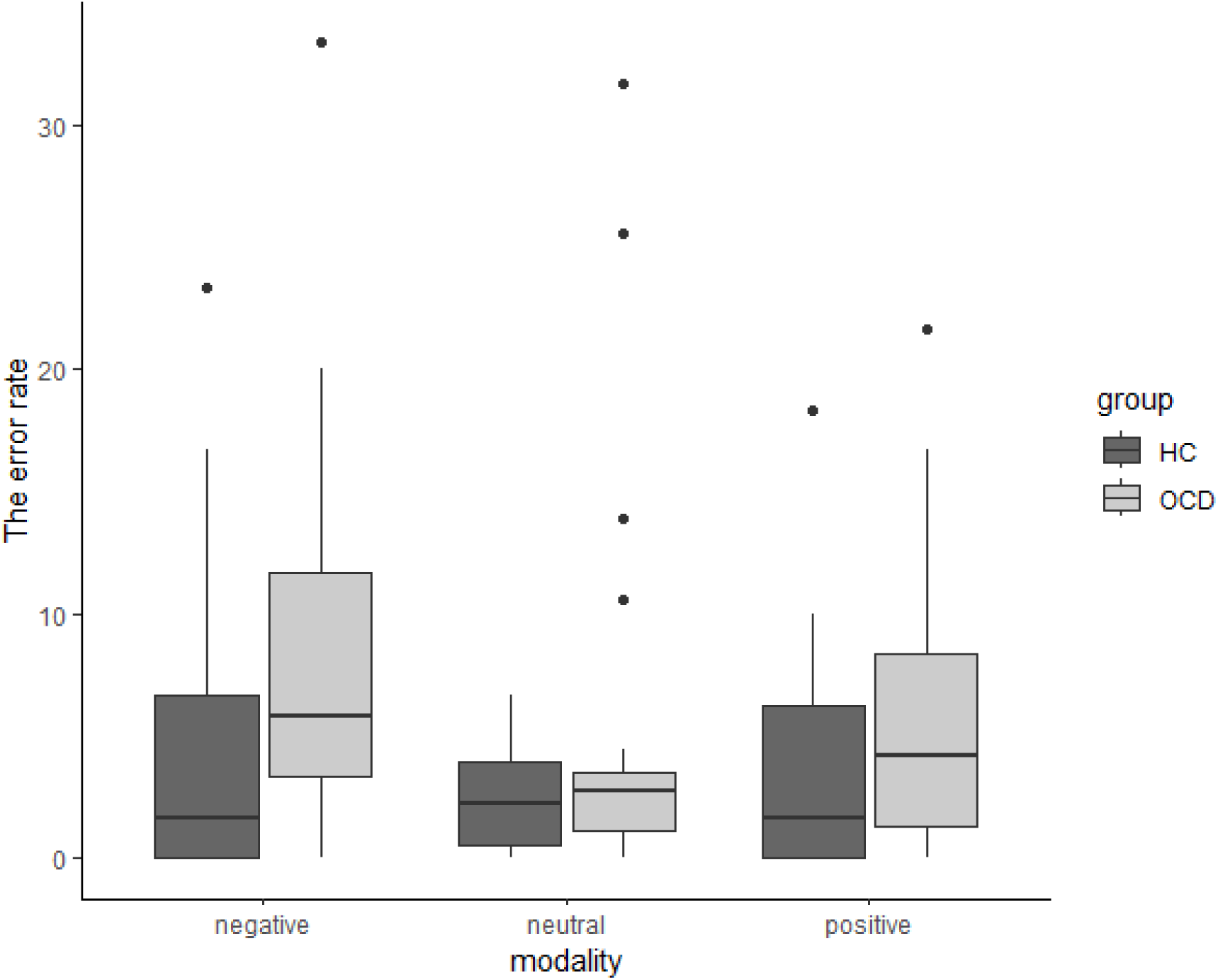
The error rate differences between groups (*HC - healthy control, OCD - obsessive-compulsive disorder)

**Table 3.**
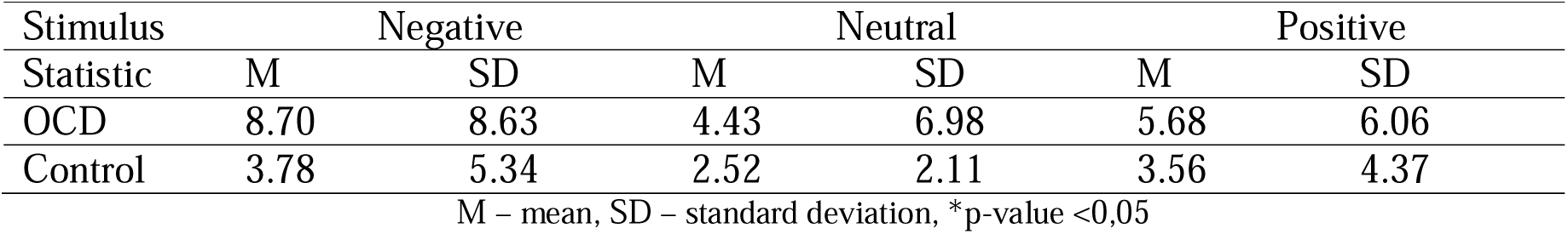
Mean and standard deviation of the antisaccade error rate.

The correct antisaccade latency (Fig.3) varied significantly between stimuli in the OCD group (p<0.01) and between valences in groups (p<0.05). The OCD group showed higher latency in negative(p<0.01) and neutral (p<0.05) stimuli than in positive stimuli. In the healthy control group, such an effect was not found. Participants with OCD made the correct antisaccade in negative and neutral stimuli with longer latency than in positive stimuli. Means and standard deviations are given in Table 4.

**Figure.3.**
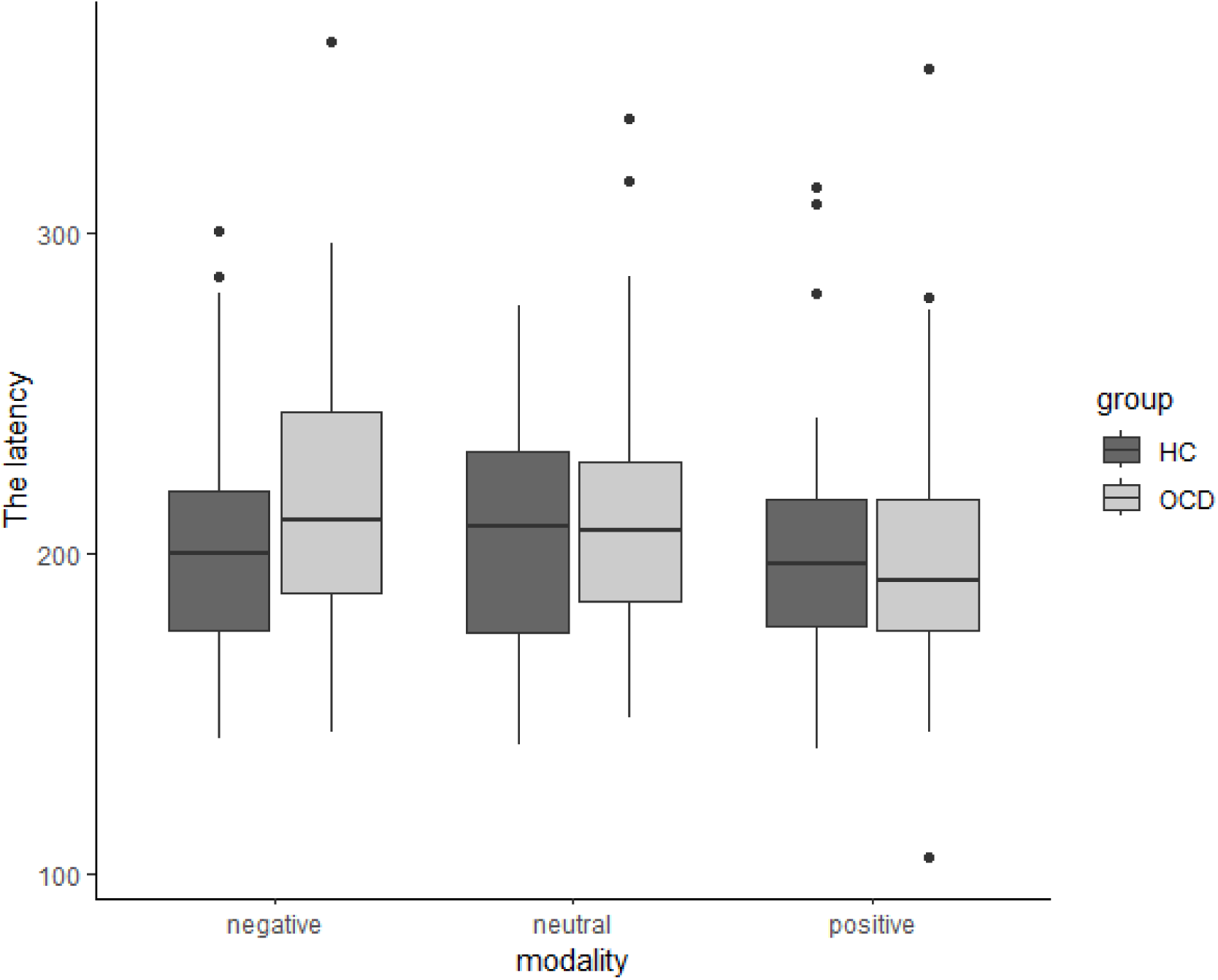
The latency differences between groups (*HC - healthy control, OCD - obsessive-compulsive disorder)

**Table 4.** Mean and standard deviation of the correct antisaccade latency.

| Stimulus | Negative |  | Neutral |  | Positive |  |
| --- | --- | --- | --- | --- | --- | --- |
| Statistic | M | SD | M | SD | M | SD |
| OCD | 219 | 47.2 | 213 | 43.6 | 199 | 47.4 |
| Control | 203 | 40.6 | 206 | 40.4 | 203 | 42.9 |

M – mean, SD – standard deviation, \*p-value < 0,05

The correct antisaccade average velocity (Fig.4) revealed differences between valences, the average velocity to neutral stimuli differed from the average velocity to positive and negative stimuli (p<0.01). Both groups made correct antisaccades faster in negative and positive stimuli than in neutral ones. Means and standard deviations are given in Table 5.

**Figure 4.**
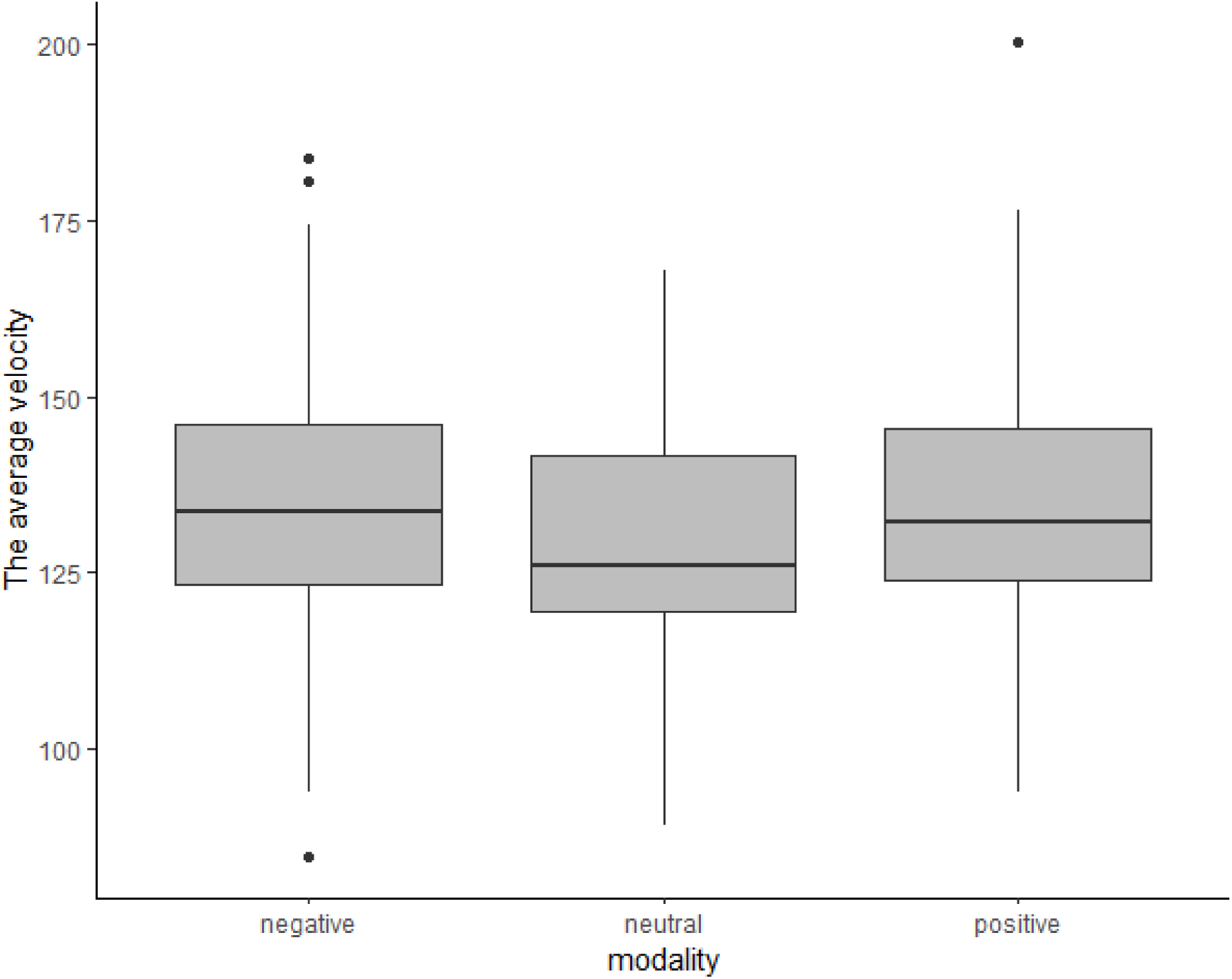
The average velocity differences between stimuli emotional valences.

**Table 5.** Mean and standard deviation of the correct antisaccade average velocity.

| Stimulus | Negative |  | Neutral |  | Positive |  |
| --- | --- | --- | --- | --- | --- | --- |
| Statistic | M | SD | M | SD | M | SD |
| OCD/Control | 135 | 19.2 | 129 | 17.1 | 135 | 19.5 |

M – mean, SD – standard deviation, \*p-value <0,05

The anticipatory saccades (Fig.5) showed differences between emotional valences (p<0.01) but did not find differences between groups. The number of anticipatory saccades was calculated in the “Stimulus - Black Screen” interval with a limit of up to 80 ms. Both groups performed more anticipatory saccades in the Stimulus - Black Screen interval to neutral stimuli than to positive and negative ones (p<0.01). Means and standard deviations are given in Table 6.

**Figure 5.**
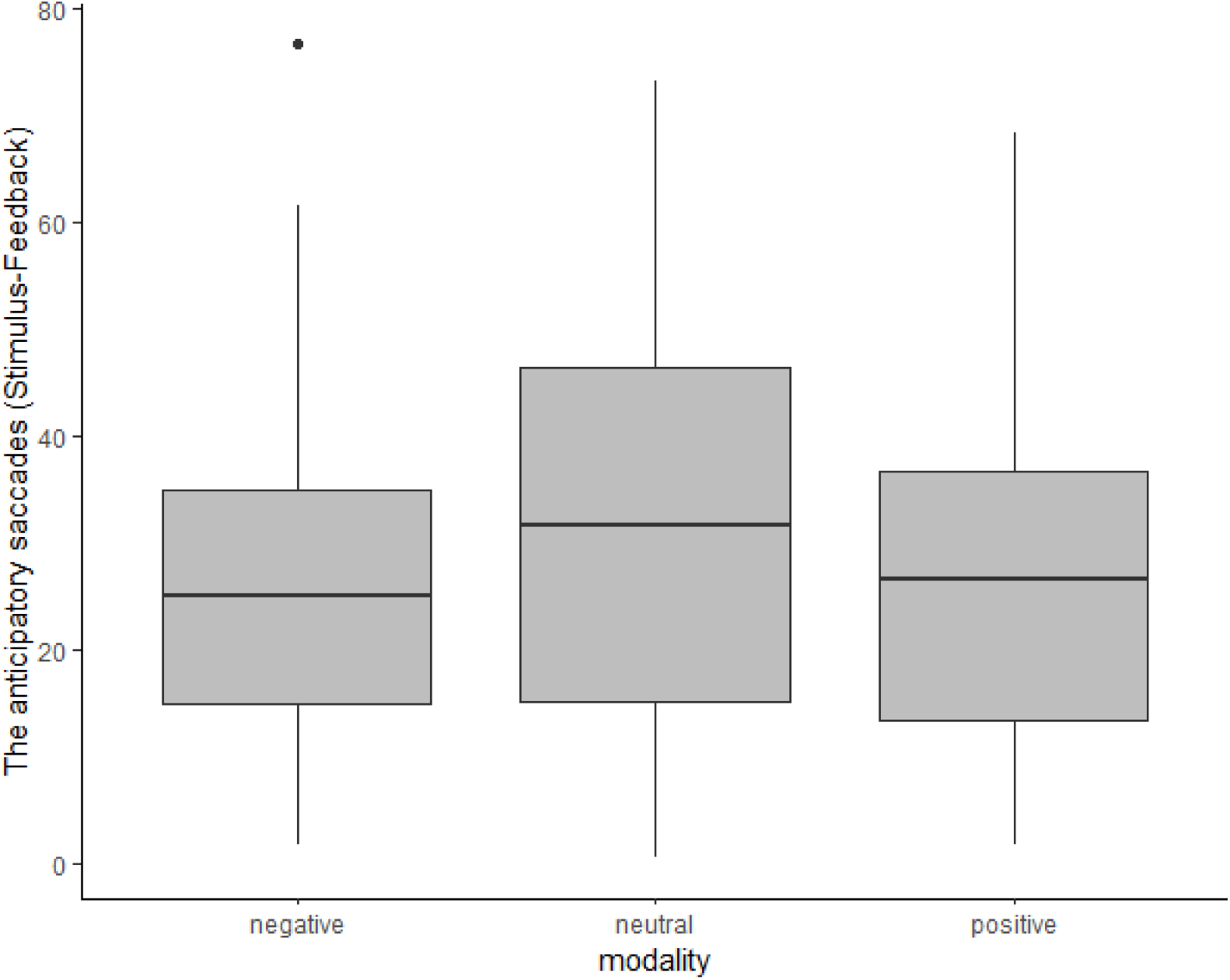
The anticipatory saccades differences in Stimulus-Black Screen interval between stimuli emotional valences.

**Table 6.** Mean and standard deviation of the anticipatory saccades (Stimulus-Black Screen interval)

| Stimulus | Negative |  | Neutral |  | Positive |  |
| --- | --- | --- | --- | --- | --- | --- |
| Statistic | M | SD | M | SD | M | SD |
| OCD/Control | 26.9 | 15.9 | 31.3 | 19.3 | 26.6 | 16.3 |

M – mean, SD – standard deviation, \* $p$ -value $< 0,05$

The peak velocity parameter did not show any differences either between groups or between valences.

## 4. Discussion

The current study used an antisaccade paradigm with multimodal stimuli (negative, positive, and neutral) to fully assess the features of cognitive and emotional control in patients with OCD. Successful execution of an antisaccade task requires two processes: suppression of the reflex saccade towards the appearance of a peripheral stimulus and voluntary eye movement to a point that mirrors the peripheral stimulus in the visual space. Typical indicators in the antisaccade task are errors, namely saccades in the wrong direction (reflecting problems of inhibitory control: the inability to suppress an inadequate visual response is most likely due to a lack of top-down control) and latency (Fischer et al., 1997). The antisaccade task measures the features of voluntary behavior and is also sensitive to dysfunction of the cortico-striatal-thalamo-cortical (CSTC) circuit (Hutton & Ettinger, 2006). Such dysfunction is the main hypothesis for the neurobiological mechanisms of OCD (Narayanaswamy et al., 2021). Previous studies using antisaccade tasks show a large variability in outcomes with respect to errors and latency.

Some studies have shown increased error rates and latency in OCD patients (Hu et al., 2020; Narayanaswamy et al., 2021), while others have argued that OCD patients do not have these indicators increased (Agam et al., 2014; Maruff et al., 1999; Spengler et al., 2006). Maruff P. et al. noted that patients with OCD experienced difficulty only when performing those tasks that required them to control movements based on an internal idea of the purpose of the task. When solving tasks in which the target remained visible throughout the entire test, no difference in the latency of saccades in patients with OCD and healthy controls was found (Maruff et al., 1999). This feature of the antisaccade task reflects the difficulty of using the internal representation of the saccade target. One of the latest studies by Bey et al. in the large sample (169 OCD, 183 healthy, 100 relatives of people with OCD) showed moderate performance impairments in the antisaccade task: increased antisaccade latencies, intra-subject variability (ISV) of antisaccade latencies, and antisaccade error rates in people with OCD compared with the healthy control group. Among other things, unaffected relatives of patients with OCD also showed elevated antisaccade express error rate and increased ISV of antisaccade latencies (Bey et al., 2018). In early studies, healthy relatives also showed both an increased error rate and increased latency (Kloft et al., 2013; Lennertz et al., 2012), and therefore the authors suggested that these eye-movement patterns may serve as a candidate for the OCD endophenotype. These particular patterns (increased antisaccade latency and error rate) may represent a general, OCD-specific cognitive deficit. Another study showed that the uncertainty of the target existence caused an increased overall check in people with OCD, which manifests itself in a large search time and fixation duration in visual search task (Toffolo et al., 2016). Uncertainty intolerance is likely to broadly influence their OCD symptoms and anxiety regardless of the manifestation of their primary symptoms (Pinciotti et al., 2021).

Interestingly, all of the aforementioned studies used a neutral stimulus as a circle or square. However, OCD pathogenetically manifests itself in the inability to stop irrelevant thoughts and actions based on emotional dysregulation (See et al., 2022). Moreover, studies targeting symptom provocation and emotion processing have found structural changes and altered functional activity in the limbic regions including the amygdala (Hazari et al., 2019), which support emotion dysregulation in these patients. Thus, we constructed an antisaccade paradigm with multimodal stimuli (negative, positive and neutral) to map out a complete picture of the psychophysiological characteristics of patients with OCD by studying oculomotor reactions.

Our finding of an increased error rate to negative stimuli in OCD patients compared to healthy controls suggests that the effect of activated threat on inhibition may play a role in the development and maintenance of OCD. Given the fact that Bannon S. et al. (2008) studied the processes of inhibition and the influence of threat on this mechanism and determined decreased inhibition under the influence of threat in patients with OCD (Bannon et al., 2008). It can be assumed that people with OCD suffer from impaired control of emotions when exposed to negative environmental stimuli.

Moreover, our results also show that latency in response to negative and neutral stimuli was significantly higher than to positive stimuli in OCD group. Current cognitive models of OCD argue that the disorder develops and is maintained through an overestimation of both personal responsibility and the level of threat posed by situations and experiences (Rachman, 1997; Salkovskis, 1999). In people with OCD, it is their "catastrophic" evaluation of obsessions and the actions they take to eliminate the negative feelings associated with them that causes their obsessive-compulsive symptoms (Frost et al., 2002; Salkovskis, 2007). Based on empirical data (Lavy et al., 1994; McNally et al., 1992), it can be concluded that participants with OCD are biased toward threatening stimuli and are slower in completing tasks with emotional negative stimuli. For example, participants with compulsive washing are slower to respond to wash words than healthy participants (Foa et al., 1993). We hypothesize that obsessive-compulsive triggers may affect the quality of attention to relevant information and the ability to suppress irrelevant stimuli. Moreover, this is also supported by Bradley’s study (Bradley et al., 2016), which proved that attentional bias is related to the theory of “delayed disengagement/maintenance” (Georgiou et al., 2005), according to which individuals with OCD are overly fixated on pictures of OCD-provoking stimuli in the later stages of processing. That supported our data that individuals with OCD have more latency of correct antisaccades in reaction to negative pictures after they are fixated on them. Another study found that the OCD group has a ‘lowered perceptual threshold’ for identifying and reacting to OCD-related material, namely it showed increased vigilance for threat during free viewing and visual search, and difficulty in disengaging from threat in visual search tasks. Participants with the high level of contamination fear showed shorter fixations on contamination threat than participants with the low level of contamination fear in the free viewing task (Armstrong et al., 2012; Armstrong & Olatunji, 2012). Various outcomes of attentional biases in eye movement responses have been found using different goals and paradigm constructs. A recent systematic review and meta-analysis of eye-tracking studies found that attentional biases can support both "vigilance" and "maintenance" theories depending on task design, which, in turn, is reflected in different eye movement outcomes (Basel et al., 2023). A higher latency to neutral stimuli than to positive ones may be due to the fact that neutral pictures are monotonous, and an individual with OCD could freeze after viewing negative pictures, and therefore the duration of latency to neutral pictures increased. Positive stimuli could switch attention from pictures of negative content in connection with the emotional valence of a different spectrum. In our opinion, it is impossible to conclude that there is a violation of the general inhibitory control, since individuals with OCD did not make errors in the pictures of neutral valence, and there were no statistical differences between the groups in relation to neutral pictures.

The results of our study showed difficulties in disengagement from negative stimuli in the OCD group in which the latency of the OCD group was greater for negative stimuli than for positive. It is possible that the inhibitory response is impaired by biased attention to negative stimuli. In general, it can be assumed that attentional biases develop as a result of the activation of negative cognitive circuits, which, in turn, cause people to focus on environmental stimuli that correspond to their main fears (Salkovskis P. M. & McGuire J., 2003). It should be noted that there were no statistically significant differences between OCD and healthy controls in the latency period due to the nature of the Overlap paradigm, namely, due to the fact that participants already knew in advance (Overlap interval) where to look. However, anticipatory saccades in the “Stimulus-Black Screen” interval did not differ between groups. This may be due to the fact that saccades in the “Stimulus-Black Screen” interval were limited to 80 ms and most of the anticipatory saccades were performed earlier.

Average velocity did not show any differences between the groups. Both groups showed increased levels in the negative and positive stimuli than in the neutral valence. It confirms previous studies showing that responses to emotionally significant stimuli are faster and have a higher priority, which in general seems to have evolutionary adaptive value (Pilarczyk & Kuniecki, 2014). The peak velocity parameter did not show any differences between groups and between valences. In any case, the calculation of this parameter is incorrect in relation to people taking drugs, since taking drugs affects the change in this parameter in oculomotor reactions (Crawford et al., 1995; Green & King, 1998; Lynch et al., 1997; Reilly et al., 2008). Our study found no difference in peak velocity, possibly due to the fact that some participants with OCD were not taking medication.

## Limitations

The present study has several limitations. Firstly, the majority of participants (23) with OCD were taking medication at the time of the data collection. 9 participants with OCD who did not take drugs and had mild OCD symptoms and could therefore live without medication. However, the results of many studies of oculomotor responses under medication (benzodiazepines, first and second generation antipsychotics, antidepressants) showed no effect of these drugs on oculomotor responses, except for a decrease in peak saccade velocity (Crawford et al., 1995; Green & King, 1998; Lynch et al., 1997; Reilly et al., 2008).

Secondly, due to the fact that our sample was fairly young, we cannot assert whether the inhibitory control and attention worsen over time in general. We expect a longitudinal study or a study in an older age category of individuals with OCD to be of great interest.

Thirdly, it would be interesting to further investigate alterations in cognitive and emotional control, dividing OCD participants into separate groups depending on the type of OCD. A study using this paradigm in groups with attention deficit and hyperactivity disorder (ADHD), who are characterized by impaired inhibitory control, as well as a study of borderline personality disorder (BPD), characterized by emotional dysregulation, can help to distinguish features of eye-movement patterns in these disorders (OCD, ADHD, and BPD).

## Conclusion

Our results imply that attentional biases toward threatening stimuli primarily affects inhibitory control in OCD. It may depend on a person’s prior experience (childhood upbringing and psychotrauma) (Salkovskis P. M. & McGuire J., 2003), which serves as a trigger that starts OCD and following difficulties in emotional dysregulation (See et al., 2022). In this study, the attentional bias is reflected in the increased latency of correct antisaccades when the fixation stimulus was a negative one in the OCD group coupled with the elevated error rate compared with the healthy controls. Many studies showed abnormal fear extinction in OCD (Cooper & Dunsmoor, 2021; Steuber & McGuire, 2022). The attentional bias to negative stimuli may serve as a fixation of cognitive scheme-beliefs which commence a vicious circle of OCD. The antisaccade task with different emotional stimuli (neutral, positive, and negative ones) can become a reliable tool for studying mental disorders based on impaired inhibitory control, attentional biases and emotional dysregulation.

The article is an output of a research project implemented as part of the Basic Research Program at the National Research Program at the National Research University Higher School of Economics.

